# Implication of exofacial thiol groups in the reducing activity of *Listeria monocytogenes*

**DOI:** 10.1101/353409

**Authors:** Guillaume Pillot, Anne Brillet-Viel, Hervé Prévost

## Abstract

*Listeria monocytogenes* growing in BHI in anaerobic condition leads a decrease of redox potential at pH7 (*E*_h7_) from 0 to −250 mV. An investigation of mechanisms involved in this reducing activity shows the implication of thiol groups located on the bacterial cell surface. Indeed, after reduction of media to −250 mV, only thiol-reactive reagents could restore the initial *E*_h7_ value. Moreover, the growth of *L. monocytogenes* in anaerobic condition is characterized by a reduction then acidification phases. This suggests a sensor mechanism of environmental *E*_h7_ to convert the metabolism way from the first phase to the second. Finally, the concentration of exofacial thiol still increase strongly during the acidification phase of *L. monocytogenes*, even after reach the minimal value of *E*_h7_. These results suggest that maintaining the exofacial thiol (−SH) groups in a reduced state do not depend on an active mechanism. Thiol groups appear to be displayed by membrane proteins or cell wallbound proteins and may participate in protecting cells against oxidative stress.

## Introduction

The oxido-reduction potential (*E*_h_) was shown to have different effects on microorganisms. *E*_h_ could be involved in gene expression, metabolism, physiology and so, could modify their growth capacity and end product formation.

The *E*_h_ measurement and monitoring are increasingly used as tools for the determination of bacterial activity in liquid media (Küsel and Dorsch, 2000), food products (Abraham et al., 2013; Alwazeer et al., 2003; Aubert et al., 2002), soils (Fiedler et al., 2007) and sediments (Schulz, 2000). Recent studies have emphasized that microbial cultures can generate a reducing activity during their growth depending on the redox potential of different cellular compartments, and their ability to grow in the presence of dioxygen. (Prévost and Brillet-Viel, 2013).

However, the mechanisms involved in bacterial reducing capacities remained poorly understood. Two mechanisms have been described. The first correspond of the consumption of oxidant compounds or the release of reducing end-products during the energy generation metabolism. Indeed, to regenerate ATP, organic substrates are oxidized and dehydrogenated during the glycolysis and citric acid cycles producing reduced electron carriers (NADH, FADH_2_). Then, these carriers are re-oxidized by electron transfer in the case of respiratory metabolism to oxidants or to metabolic intermediates in fermentative metabolism. Thus, the metabolic pathways of energy production determine the reducing activity of microorganism to whether they are aerobic, facultative anaerobic or obligate anaerobic. Indeed, the terminal electron acceptor in respiration of strict aerobic bacteria is dioxygen, which restricts the range of redox potentials to values close to the oxidant values. Anaerobes meanwhile can reduce external terminal electron acceptors such as NO_3_^−^, SO_4_^2−^, Mn (III / IV) and Fe(III) which allow higher reducing capacities (until *E*_h_ = −600mV). Finally, most of anaerobes can also produce strong reducing end-product such as H_2_ (midpoint oxidation reduction potential, *E*_0_’ = −420 mV).

The second mechanism involved in bacterial reducing capacities which appears to be not linked to metabolism (proton motive force) or production of reducing end-product have been recently described in *Lactococcus lactis.* In this lactic acid bacteria, the reducing activity is due to thiol groups of cell surface proteins (Michelon et al., 2010).

The aim of this study was to identify the mechanism involved in the reducing activity of *Listeria monocytogenes.* Indeed, *L. monocytogenes* is a well-know food born pathogen with a high reducing activity. It is able to decrease the *E*_h7_ (*E*_h_ at pH 7) to −250 mV (Ignatova et al., 2008, 2010). However, the mechanism of this reducing activity was never studied

## Materials and Methods

### Bacterial strains, culture conditions

Three strains of *L. monocytogenes* were used in the present study. *L. monocytogenes* CIP 78.35 and EGDe (CIP 107776), initially isolated from veterinary and medical case with serotype 3 b and 1/2a respectively (Comi et al., 1997), were from the collection of Pasteur Institute of Paris (France) and SOR100, isolated from industrial process with unkown serotype, from the collection of LUBEM (Université de Bretagne Occidentale, France). Concentrated stock cell suspensions were stored at −80°C in Brain Heart Infusion (BHI, AES, France) supplemented with glycerol (10% v/v). Preculture of *L. monocytogenes* used as inocula were grown aerobically in 10 mL BHI broth during 12h at 37°C. As need, *L. monocytogenes* was enumerated on BHI agar plates incubated at 37°C during 24h.

### Redox potential and pH measurement

To prevent any uncontrolled *E*_h_ modification of the BHI broth due to reaction of Maillard during autoclaving, BHI media was sterilized by filtration at 0.2 μm as described by Ignatova et al. (2009). Anaerobic condition was reached by nitrogen bubbling (Alphagaz 1, Air Liquide, France) during 5 hours. Then, BHI media was inoculated with 1 × 10^5^ cells.mL^−1^. All growth and reducing activity experiments were conducted in a 400-ml water-jacketed double-sidearm glass bioreactor (designed in Batailler labo, La Chapelle sur Erdre, France) especially customized with butyl rubber seal caps (Laboratoires Humeau, La Chapelle sur Erdre, France) to ensure air tightness. The redox potential (*E*_m_) and pH of culture were monitored at 37°C on BHI using a combined redox and pH electrode (InPro 3253SG/120/PT100, Mettler Toledo, Viroflay, France). The redox electrode was referred to the Ag/AgCl system. The electrode was connected on-line to a multi-channel interface (Absciss interface; Absciss, Fixin, France). In order to obtain better stabilization of measured redox potential, the electrode was polished with aluminum powder before each experiment and treated with HCl- and pepsin-based Electrode Cleaning Solution (Grosseron, Nantes, France) for 30 min once per week. A control measure was performed in tap water before each experiment (Abraham et al., 2013). The measured redox values collected via the interface were converted to *E*_h_ and *E*_h7_ values by using the software “Multi-bioreactors Central (3x *E*_h_, 3xpH, 3xT°C)” (Absciss, Fixin, France) using the equation *E*_h_ = *E*_m_ + *E*_ref_. In this equation *E*_h_ is the redox potential compared to the standard hydrogen electrode, *E*_m_ the measured redox potential and *E*_ref_ the standard potential of the reference electrode. The standard potential of the reference Ag/AgCl electrode used in this study, was *E*_ref_ = 196.6 mV at 37°C. To overcome the Nernstian effect of pH on the *E*_h_ value, the *E*_h_ was corrected to *E*_h7_. The *E*_h7_ which correspond to *E*_h_ if measured at pH 7 was calculated using the Leisner and Mirna equation, *E*_h7_ = *E*_h_ − [α(7 − pH)] (Leistner and Mirna, 1959) where α is the correlation factor between pH and *E*_h_. The α correlation factor for BHI at 37°C determined in this study is identical to the previously published α value for this medium (35 mVpH.U^−1^, Ignatova et al., 2010).

### Filtration

*L. monocytogenes* cells were harvested from culture by filtration using polyethersulfone 0.22 μm Steritop-GP Filter Units (Millipore, Carrigtwohill, Ireland). Polyethersulfone membrane filters were used because to their low protein and drug binding properties. The resulting filtrate was degased using nitrogen bubbling during 3h before *E*_h_ measurement.

### Neutralisation of thiol groups

In order to determine whether thiol-containing proteins and polypeptide chains are exposed at the extracellular or cytoplasmic membrane surface, two thiol-reactive reagents, the N-Ethylmaleimide (NEM) and the *4-acetamido-*4′*-maleimidylstilbene-*2,2′- disulfonic acid (AMdiS) were used in this study. NEM is a membrane permeable compound able to react with intra- and extra-cellular thiol groups. AMdiS is a high polarity water soluble compound unable to cross the membrane and then can only react with the extracellular thiol groups.

NEM were obtained from Sigma (E3876, Sigma Aldrich, St Quentin Fallavier, France) and AMdiS from Invitrogen (A-485, Invitrogen, Carlsbad, USA). A stock solution of 1M NEM was prepared in methanol: water (3 : 1). The NEM was added to a final concentration of 25 mM in culture media. A stock solution of 65.2 mM AMdiS was prepared in water and added in culture media to a 9 mM final concentration.

### Titration of accessible exofacial thiol groups

The Ellman’s method (Ellman and Lysko, 1979) using the 5,5′-dithiobis-2-nitrobenzoic acid (DTNB) was used to assay the exofacial thiol groups during bacterial growth. Because DTNB is a membrane impermeable compound, only the thiol groups located at the bacterial cell surface could react. Thiols react with DTNB, cleaving disulfide bond to give 3-thio-6-nitrobenzoate (TNB^−^), ionized into a yellow colored TNB^2−^ dianion.

*L. monocytogenes* cells were harvested from the culture by centrifugation for 15 min at 3500g then suspended in 1mL of 0,1 M potassium phosphate buffer, 60 μM DTNB, (pH 8). The cell suspension was incubated 30 min at room temperature in the dark then centrifuged for 15 min at 3500g. The supernatants were filtered through a 0.45 μm membrane filter (Millipore). The concentration of exofacial thiol groups in the filtrate was measured by spectrophotometer at A_412_ and the concentration was calculated using N-acetyl-L-cysteine standard curves (range: 5-100 μM).

### Statistical analysis

The data analysis was performed using the statistical analysis software Statgraphics Centurion XVI (StatPoint Technologies, Inc., Warrenton, Virginie, USA). An analysis of variance (ANOVA) with α ≤ 0,05 was used to compare the results.

## Results and discussion

### Reducing activity of *L. monocytogenes*

The average evolutions (eight experiments) of *E*_h7_ and pH of *Listeria monocytogenes* SOR 100, EGDe and CIP 78.35 growing at 37°C on BHI under anaerobic conditions are presented figure 1. The kinetics of *E*_h7_ and pH during growth of *L. monocytogenes* shown two consecutive phases. The first corresponds to a reduction and a constant pH phase during the 5 first hours of culture. This phase is characterised by the reduction of the media from initial *E*_h7_ value (1 to 24 mV) to a minimal *E*_h7_ (−232 to −245 mV) depending of the strain. During the reduction phase, the pH remainded stable at 7.3. After the reduction phase the *E*_h7_ remained stable until the end of the growth of *L. monocytogenes* but the pH decreased. The second phase corresponds to an acidification phase where the pH decreased from 7.3 to 5.9 between the 5^th^ hour to the 13^th^ hour of culture, then slowly to a final pH value around 5.5 at 48h.

**Fig. 1.**
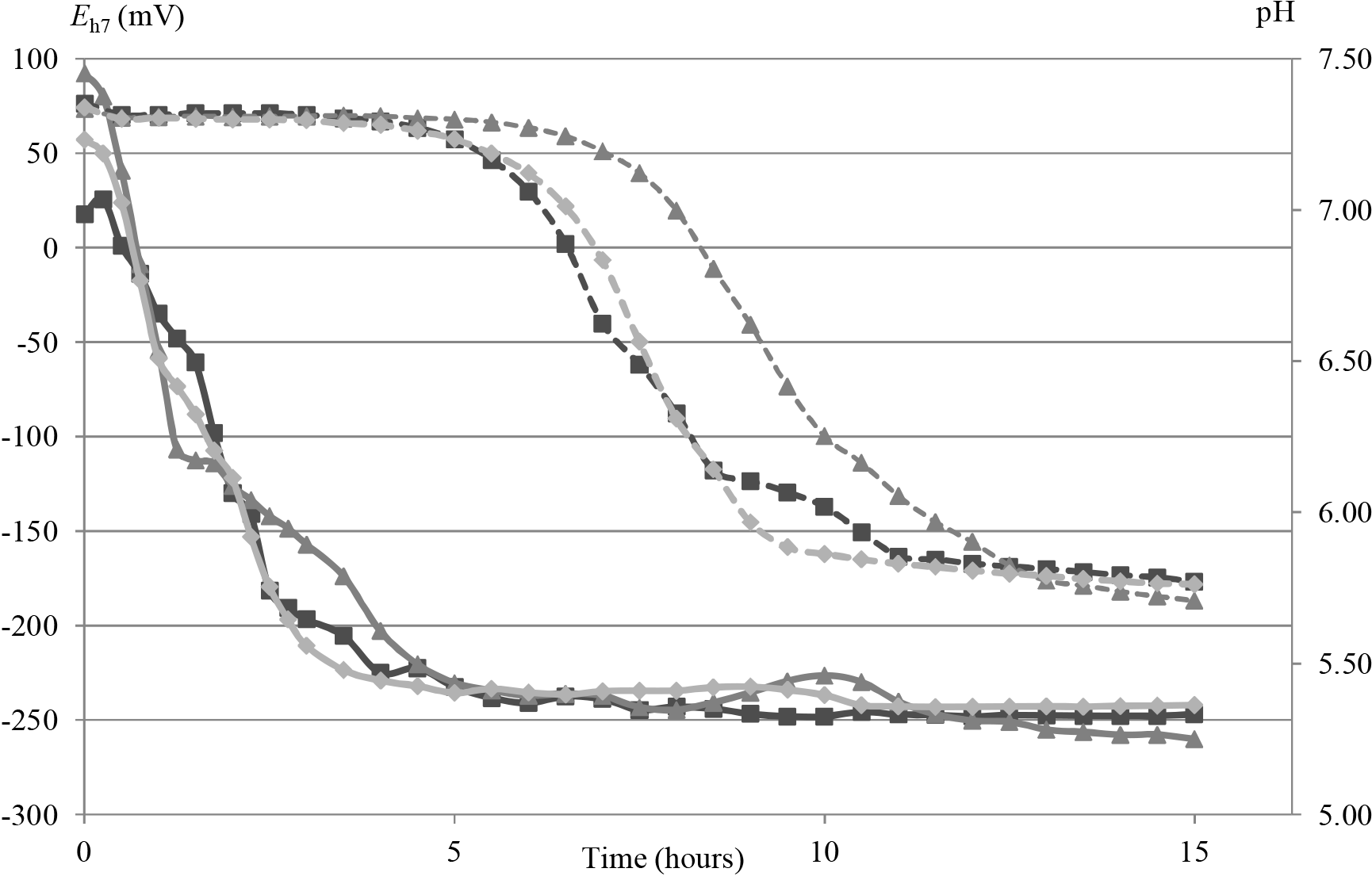
Evolution of Eh7 (—) and pH (---) during culture of *Listeria monocytogenes* SOR 100 (■), EGDe (▲) and CIP 78.35 (♦) under anaerobic conditions (N_2_). Average curves of 8 experiments are shown (SD respectively for SOR 100, EGDe and CIP 78.35 are for *E*_h7_ = ± 35 / 47 / 26 mV and pH = ± 0.17 / 0.15 / 0.10 pH units)

The descriptive factors of reduction and acidification phases for the three strains are presented Table 1. The descriptive factors from the reduction and acidification kinetics were the initial and final values of *E*_h7_ and pH, the lag time and duration of each phase and the mean rate of reduction and acidification. Any significant difference was shown between SOR 100, EGDe and CIP 78.35 by statistical analysis for each parameter.

**Table 1.** Descriptive parameters of reduction and acidification phases of the growth of *L*.*monocytogenes* and *L. lactis* in anaerobic conditions. ANOVA (P=0,05) was used to statistical analysis. Each value is an average of 8 experiments. Any difference was observed between SOR 100, EGDe and CIP 78.35.

|  | *Listeria monocytogenes* in BHI |  |  | *L. lactis ssp cremoris* |
| --- | --- | --- | --- | --- |
|  | SOR 100 | EGDe | CIP 78.35 | *TIL 46* in MRS*<br>(Michelon et al., 2010) |
| Initial Eh<sub>7</sub> (mV) | 1 ± 70 | 17 ± 92 | 24 ± 102 | 148 ± 19 |
| Lag time of reduction (h) | 0.7 ± 1.03 | 0.75 ± 1.47 | 0.53 ± 0.57 | 1.84 ± 0.1 |
| Rate of reduction (mV.h<sup>-1</sup>) | -96 ± 57 | -90 ± 88 | -100 ± 95 | -130 |
| Time of reduction (h) | 3.6 ± 1.1 | 4 ± 1 | 4.3 ± 1.5 |  |
| Final Eh<sub>7</sub> (mV) | -245 ± 23 | -232 ± 34 | -233 ± 21 | -219 ± 1 |
| initial pH | 7.31 ± 0.03 | 7.31 ± 0.04 | 7.34 ± 0.05 | 6.37 ± 0.06 |
| Lag time of acidification (h) | 4.9 ± 1.5 | 5.5 ± 1.02 | 6.6 ± 2.9 | 3.43 ± 0.04 |
| Rate of acidification (pH units.h<sup>-1</sup>) | -0.36 ± 0.27 | -0.38 ± 0.18 | -0.38 ± 0.21 | -0.13 |
| Time of acidification (h) | 4.31 ± 2.75 | 4.5 ± 2.2 | 4.51 ± 2.52 |  |
| Final pH (pH units) | 5.88 ± 0.07 | 5.86 ± 0.08 | 5.90 ± 0.09 | 5.97 ± 0.06 |
| pH at 48h (pH units) | 5.54 ± 0.14 | 5.49 ± 0.07 | 5.69 ± 0.07 | 4.57 ± 0.06 |

To date, the reducing activity of *L. monocytogenes* has been only studied in aerobic conditions (Ignatova et al., 2010). In fact, in aerobic condition, the reduction and acidification are not divided into two separated phases and occur in the same time. Moreover, the final value of *E*_h7_ is different between the two atmospheres, with a value of −179 ± 38 mV in aerobic condition against −245 ± 23 mV in anaerobic condition both independently of the culture medium and the initial *E*_h7_ value. This difference of minimal *E*_h7_ value is certainly due to the presence for the first case of dissolved dioxygen in the media, known to be a high oxidative compound. In addition, the only study on the reducing activity of another bacterium, *L. lactis*, allow to compare our results to this Lactic Acid Bacteria (Michelon et al., 2010). Comparatively, the reduction phase of *L. lactis* occur later than *L. monocytogenes* with a *E*_h7_ value higher, while the acidification phase starts sooner with a pH value at 48h lower.

### Role of the cell in the E_h_ decrease

In order to study the cell-linked reducing activity, the *E*_h7_ of *L. monocytogenes* cultures at the end of the reducing phase (ERP), cell-free ERP filtrate and sterile BHI were compared (Figure 2). The *E*_h7_ value measured in sterile BHI were range from 10 to −40 mV. The *L. monocytogenes* culture reduced the medium to −240 mV at ERP. The statistical analysis show that for the three *L. monocytogenes* strains, *E*_h7_ of cell-free ERP filtrate and sterile BHI are not significally different (α ≤ 0,05). These results shown (i) that reducing activity of *L. monocytogenes* is not due to production of reducing end-products from metabolism released in the medium (ii) that the restoration to the initial *E*_h7_ results from the cell removing from the medium. So, the reducing activity of *L. monocytogenes* culture is mediated by the whole cells. A same observation had been made on *L. lactis* where the reducing activity was mediated by the cells (Michelon et al., 2010).

### Implication of Thiol groups in the decrease in E_h_

Having refuted the hypothesis of reducing molecules excreted by the cell, we have investigated the thiol cellular mechanism. To study the role of thiol compounds, two reagents, N-Ethylmaleimide (NEM) and 4-acetamido-4’ -maleimidylstilbene-2,2’-disulfonic acid (AMdiS) were selected. These reagents contain maleimide, which can bind with thiol groups in an irreversible reaction to suppress there activity in redox equilibrium. Due to its neutrality, NEM can penetrate the cytoplasmic membrane and neutralized intra-and extra-cellular thiol compounds. Contrariwise, AMdiS which is negatively charged cannot diffuse across the cellular membrane. Thus, AMdiS can only react with external thiol groups. As shown in the Figure 2, the addition of NEM or AMdiS to a media reduced by *L. monocytogenes* increase the *E*_h7_ to its initial value. Therefore the experiment with NEM show that thiol groups are engaged in reducing activity, while AMdiS indicate their exofacial localization. Thus, we can assume that these ERP values are likely the result of an equilibrium of thiol redox compounds as glutathione (*E*_0_’ = −240 mV), thioredoxin (*E*_0_’ = −195 to −270 mV) or protein carrying cysteine (*E*_0_’ = −340 mV) (Mössner et al., 1999; Schafer et al., 2001; Sober et al., 1968). Bioinformatics sequence analysis of *Listeria monocytogenes* EGD-e has identified five proteins with a thioredoxin CXXC motif, may play a role in reducing mechanisms. Among these proteins, there are four proteins with a T-X-X-C pattern and one with a C-X-X-T pattern. Analysis of these proteins have identified as a thiol peroxidase, a peroxiredoxin, a ribonucleotide reductase and glutathione peroxidase.

**Fig. 2.**
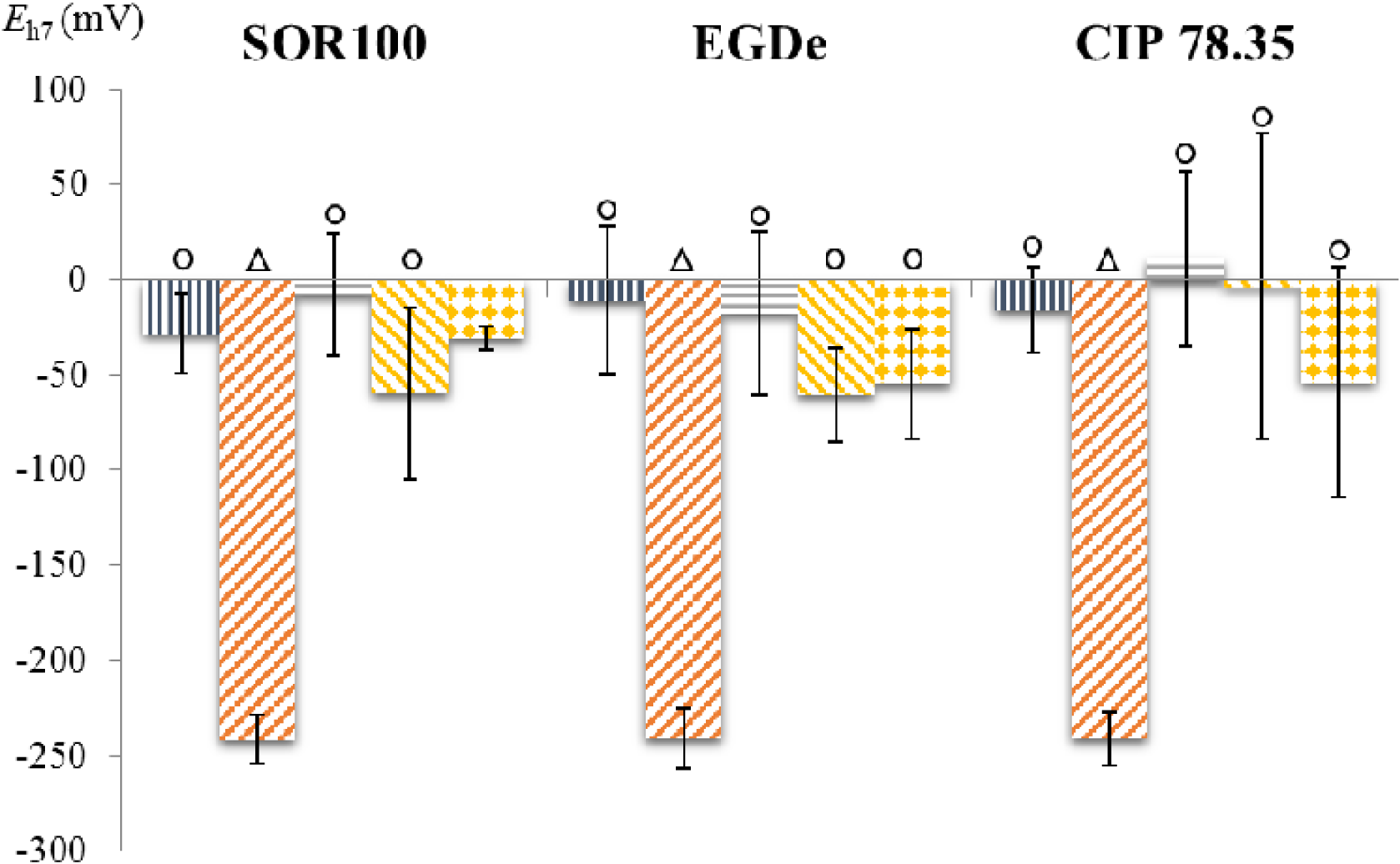
Histogram of *E*_h7_ values before inoculation, after reduction, after filtration, and after addition of NEM or AMdiS in culture, for *Listeria monocytogenes SOR 100, EGDe* and *CIP 78.35*. For each treatment, the colums shown the average of a triplicate. The statistical analysis was performed with ANOVA (P=0.05) and significant difference are show by ◯ and △ above the columns.

### Evolution of exofacial thiol groups concentration during growth of *L. monocytogenes*

The evolution of cell and exofacial thiol concentrations during the culture of *L. monocytogenes* SOR 100, EGDe and CIP 78.35 in anaerobic BHI media are represented Fig 3A. The bacterial population and exofacial thiol concentration increase simultaneous exponentially. Indeed, during the reduction phase, the concentration of exofacial thiols increase slowly from 0 to 20 ±5 μM while during the acidification phase the concentration increases to a maximum of 82 ±22 μM. Thus, the concentration of exofacial thiol still increases after the end of reduction phase, unlike *L. lactis* (Michelon et al., 2010) where the concentration stays stable at the end of reduction at 12 μM until the end of growth.

**Fig. 3.**
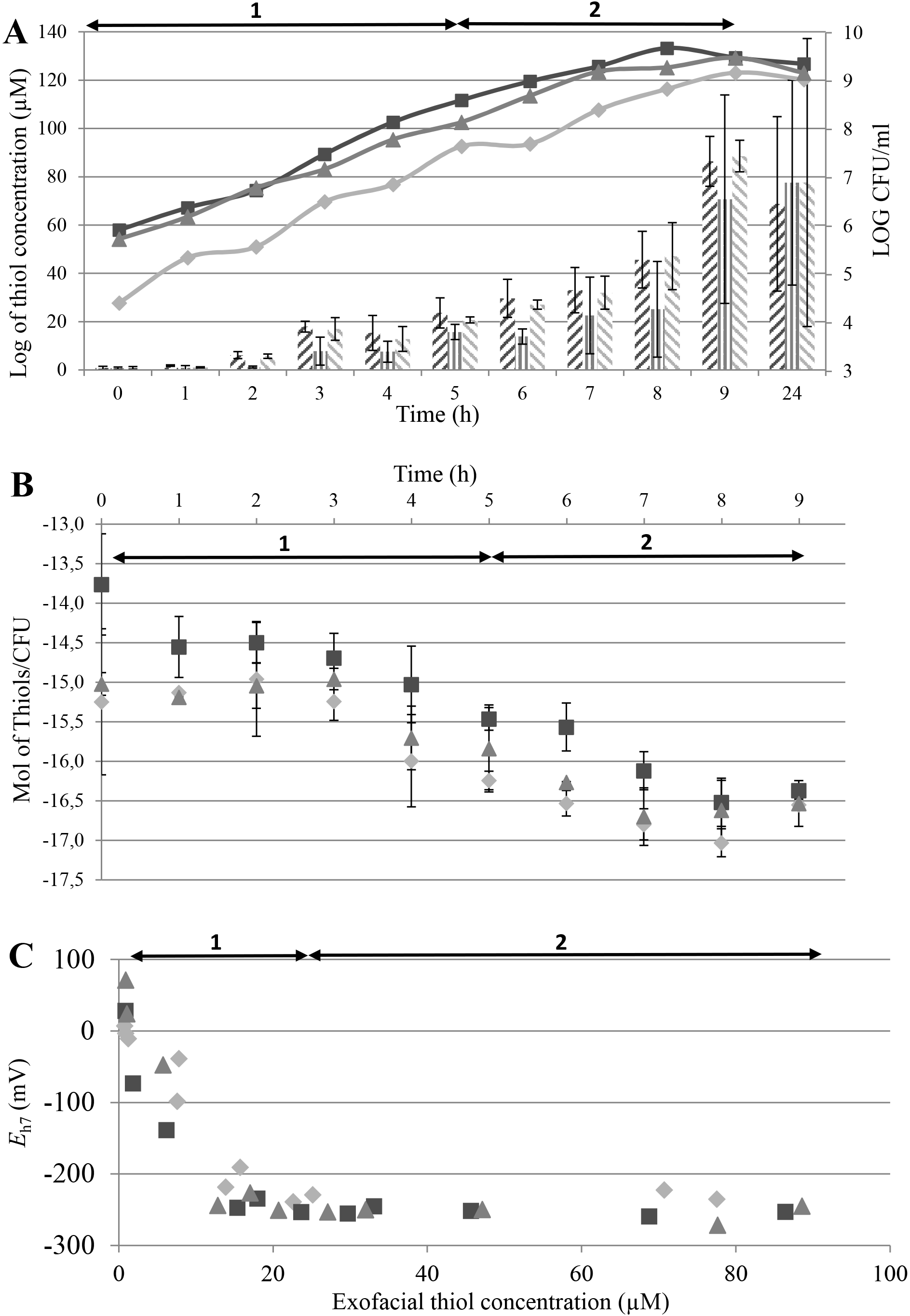
Evolution in the concentration of exofacial thiol groups during reduction by *L. monocytogenes SOR 100* (■), EGDe (▲) and CIP 78.35 (♦) under anaerobic conditions (N_2_).(A) Evolution of exofacial thiol concentration (histograms) and growth (curves). (B) Evolution of the rate of mole of exofacial thiol per CFU. (C) *E*_h7_ evolution depending on the exofacial thiol concentration. Phase 1 is the reduction phase while Phase 2 is the acidification phase.

However, the ratio of thiol concentration per cell (Fig. 3B) is stable during the reduction phase with a value of 1 to 5 femtomol/CFU and decrease during the acidification phase to a final value of 30 attomol/CFU. Thus, the ratio is divided by a 100 factor during the acidification phase.

The fig 3C shows the correlation between thiol concentration and *E*_h7_ value of the media. During the reduction phase, the *E*_h7_ of media decrease proportionally to the increase of exofacial thiol concentration. However, when the value of 20 μM of thiol is reached, the *E*_h7_ of media stay stable at a minimum of −250mV while the thiol concentration still increases to a range from 70 to 88 μM at 9h. This suggests that the Eh was stabilized when the concentration of exofacial thiol molecules in the media reach a limit value (≈ 20 μM). From this concentration the thiol–disulfide redox couple establish a more intense current with the Pt electrode than other redox couples in the culture medium (Michelon et al., 2010; Peiffer et al., 1992).

In contrast with *L. lactis* (Michelon et al., 2010), the concentration of exofacial thiol of *L. monocytogenes* still rises strongly until the stationary phase, while the ratio of thiol per cell decrease. These results suppose a conversion of the metabolic ways to reduce the production of exofacial thiol when the limit value is reached. Although the reducing activity was not directly related to metabolic activity, reducing equivalents such as NADH or thioredoxin are likely to be involved in the formation of exofacial thiol groups during the reducing phase. These results let us suppose a sensor mechanism inducing a decrease of thiol compounds production when a minimum of environmental *E*_h7_ value is reached. This implies a sensor mechanism inducing the modification in the gene expression of the metabolism. In fact, it’s known that bacteria are able to sense the extra or the intracellular environmental redox state with redox sensing mechanisms related to the thiol–disulfide balance and adapt their cell activity (Green and Paget, 2004). Williams et al., (2005) have shown the sequence of 2 sensor genes of environmental stress in *Listeria monocytogenes*, Lmo1948/Lmo1947, homolog of a ResDE system of *Bacillus subtilis.* This latter is composed of a membrane sensor and a cytoplasmic regulator and is involved in the regulation of aerobic and anaerobic metabolism (Duport et al., 2006; Zigha et al., 2007). In addition, another study showed a gene family of Crp / Fnr and their importance in resistance to oxidative stress by *Listeria* mutation targeted genes (Lampidis et al., 1994; Uhlich et al., 2006). The identification of proteins located on the extracellular surface and involved in the sensor mechanism and the decrease of *E*_h_ would be of interest for increasing our understanding of the mechanisms involved as well as the reducing activity of *L. monocytogenes*. Indeed, the systematic reduction of the extracellular microenvironment may participate in protecting cells against oxidative stress and others stress. Indeed, an overexpression of trxB gene encoding the thioredoxin reductase have been shown in *Listeria monocytogenes* EGD-e at low temperature growth (Liu et al., 2002). Another recent study demonstrated the presence and synthesis of glutathione in *Listeria monocytogenes* following stress caused by gallic acid and nisin, both molecules with antimicrobial activities (Moro et al., 2012). Finally, it could also be able to play a role in some demonstrated thiol-dependent mechanisms, as the folding of exoproteins or the virulence by the action of listeriolysin (Leimeister-Wachter and Chakraborty, 1989).

## Conclusions

In conclusion, the present study have shown the role of exofacial thiol in the reducing activity of *L. monocytogenes* in anaerobiosis and ruled out the implication of reducing end-products or the consumption of oxidizing compounds, as is mainly observed for other bacterial species. The decrease of *E*_h_ during the reducing activity was directly connected to the increase of cell density and concentration of thiol groups carrying on proteins on the bacterial cell surface. Exofacial thiol could establish a reducing microenvironment around the cell and might be used as ligands to coordinate such redox-responsive clusters (Green and Paget, 2004).

## Acknowledgments

This work has been done as in part of the FoodRedox research grant (ANR/ALID 2012-2015). We would like to thank all members of FoodRedox for their technical aid.

